# Effect of plant tissue culture parameters on the ploidy level of *Physalis grisea, Solanum lycopersicum*, and *Solanum prinophyllum* regenerants

**DOI:** 10.1101/2025.05.01.651681

**Authors:** Kerry Swartwood, Yumi Green, Iacopo Gentile, Zachary B. Lippman, Joyce Van Eck

## Abstract

Plants regenerated from seedling explants (hypocotyls and cotyledons) of the Solanaceae family members *Physalis grisea* (groundcherry), *Solanum lycopersicum* (tomato), and *Solanum prinophyllum* (forest nightshade) were used to determine the in vitro culture parameters that contribute to the incidence in polyploidization of tissue culture-derived plants (regenerants) from these species. We examined the possible effects of zeatin concentration in the plant regeneration medium, explant source, and species. Plants were grown to maturity under greenhouse conditions, pollen was collected and germinated. Flow cytometry analysis verified the utility of the pollen germination method for determining differences in ploidy, which was based on the number of pollen tubes produced with one tube representing diploid and two indicating polyploid. As for zeatin concentration, we assessed the effect of our standard method of initiation on medium containing 2 mg/l followed by 1 mg/l 2 weeks after culture initiation in comparison with 0.25, 0.5, and 1 mg/l throughout the culture lifetime. There were no major correlations for zeatin concentration on ploidy status across the species except for plants regenerated from *S. lycopersicum* hypocotyl explants where the percentage of polyploid regenerants increased with increasing concentrations. As for species and explant effects, *P. grisea* plants regenerated from hypocotyl explants had the highest percentage of polyploid plants at 81% compared to 43% and 35% for *S. lycopersicum* and *S. prinophyllum*, respectively. From cotyledons, 8% of *S. lycopersicum* and 20% of *S. prinophyllum* were polyploid. A comparison with *P. grisea* could not be made because cotyledon explants do not regenerate on zeatin-containing medium. The results indicated the incidence of polyploidization cannot be generalized for zeatin concentration, however, an influence of explant type and species was observed. Effects of increased ploidy on plant morphology were primarily larger flower and seed size; however, no significant differences were observed in plant or fruit size.

## Introduction

Plant tissue culture is a fundamental technique that involves the in vitro cultivation of cells, tissues, and organs on specialized medium containing components such as salts, vitamins, growth regulators, and sucrose. It has varied applications in plant breeding, genetic engineering, genome editing, conservation, and plant propagation. While plant tissue culture plays a vital role in these applications, there is a potential unintended outcome, somaclonal variation, that can lead to spontaneous changes in chromosome structure and number that sometimes causes regenerated plants to be either polyploid or aneuploid [1]. Variation in ploidy status or the number of complete sets of chromosomes in a plant’s cells is the predominant change that results from the tissue culture process [2, 3]. Various parameters in the process have been shown to contribute to ploidy variation, which is often induced because of rapid cell division from exposure to plant growth regulators in the medium or prolonged culture periods [4, 5]. It can also be caused by the type of explant and the ploidy status of cells within explants [6, 7]. Reports in the literature have documented the increased ploidy of plants from tissue culture of various plant species [6–10]. The changes in ploidy status of plant regenerants compared to the starting plant species have been attributed to three different genetic sources: 1) a mixture of cells with different ploidy levels in the explants, 2) endoreduplication, and 3) endomitosis [11, 12].

Polyploid plants, which have more than two sets of chromosomes, often exhibit increased cell size, leading to larger overall plant stature, flowers, and fruit compared to their diploid counterparts, which in turn can provide ecological benefits such as increased vigor potentially leading to better stress tolerance [13, 14]. On the other hand, polyploidy can also have negative consequences such as on reproductive traits, sometimes leading to sterility or shifts in flowering time, which can impact plant breeding and evolution [14]. Thus, ploidy status is a crucial factor in shaping plant morphology and adaptation.

The work described here was to determine how tissue culture factors influence the ploidy status of our solanaceous species of interest: *Physalis grisea* (groundcherry), *Solanum lycopersicum* (tomato), and *Solanum prinophyllum* (forest nightshade). *P. grisea*, is an underutilized plant species grown for its nutritious, sweet, yellow fruit [15, 16]. It is only semidomesticated, therefore it has undesirable traits that limit its wider adoption for agricultural production [17]. To correct these characteristics, a genome editing approach is being utilized to both understand the underlying genetic mechanisms of these traits while also developing strategies for their improvement [16, 18–21]. However, we observed that the flower and fruit phenotypes of *P. grisea* plants regenerated from in vitro cultured hypocotyl explants differed from wild type. The mature regenerated plants had larger flowers and fruit reminiscent of what would be expected from a polyploid version of *P. grisea*, which is a diploid species. To determine the possible cause(s) of this tissue-culture induced polyploidization and if the phenomenon was species-specific, we also evaluated regenerated plants from *S. lycopersicum, S. prinophyllum*, which are also part of our research program.

### Plant material

Seeds from *Physalis grisea, Solanum lycopersicum* cv M82, and *S. prinophyllum* were surface disinfected and cultured for germination according to methods previously reported [22, 23]. For each species, 6 - 8-day-old seedlings, at a stage before true leaves emerged, were used as a source of explants. Hypocotyl segments (0.5 – 1 cm in length) were excised from *P. grisea* seedlings and cultured on a plant regeneration medium as described by Swartwood and Van Eck [23]. Cotyledon and hypocotyl segments (0.5 – 1 cm in length) were used for plant regeneration of *S. lycopersicum* and *S. prinophyllum* according to a report for tomato [22]. Based on previous work, we found *P. grisea* cotyledons do not regenerate plants [23].

### Plant regeneration medium

We compared the ploidy level of plants regenerated according to our standard method where cotyledon and hypocotyl explants are maintained on a Murashige and Skoog (MS)-based medium containing 2 mg/l zeatin for 2 weeks followed by transfer of cultures to medium containing 1 mg/l with medium containing 0.25, 0.5 and 1 mg/l for the entire culture process [22–24]. All cultures were transferred to freshly prepared medium every 2 weeks. When regenerated shoots were approximately 2 cm tall they were transferred to rooting medium [22, 23]. Culture conditions and methods for acclimation to greenhouse conditions for all 3 species were according to earlier reports [22, 23].

### Ploidy determination

There are various methods that have been reported for determining ploidy level of plants [25]. We chose a method based on pollen germination and the number of pollen tubes [26]. Pollen germination medium (PGM) contained the components listed in Table 1. The medium was vacuum filtered sterilized, aliquoted into 25 – 30 ml portions, and stored at -20C. Pollen was collected from dehiscing flowers into 1.5 ml microfuge tubes to which 1 ml of PGM was added followed by vortexing for 30 – 60 sec. The tubes were placed in a centrifuge and spun down for 1 min at 5000 rpm. The pollen was resuspended and transferred into wells of either a 12- or 24-well plate. Additional PGM was added to dilute the preparation in a well if the solution was turbid, which indicated there was a large amount of pollen collected. The plate was covered with aluminum foil and incubated on a rotary shaker at 28C, 100 rpm for 2 – 4 hrs before visualizing under a stereomicroscope for pollen tube counts

**Table 1:** Pollen germination medium (PGM).

| Component | Stock solution concentration | Stock solution per liter (ml) | Final concentration |
| --- | --- | --- | --- |
| PEG 4000 | 40% | 600 | 60 mM |
| Boric Acid | 0.1% | 100 | 1.6 mM |
| Sucrose | 40% | 50 | 58.4 mM |
| HEPES buffer | 0.5M, pH 6.0 | 40 | 20 mM |
| Ca(NO<sub>3</sub>)<sub>2</sub>·4 H<sub>2</sub>O | 0.1M | 30 | 3 mM |
| MgSO<sub>4</sub>·7H<sub>2</sub>O | 2% | 10 | 756 uM |
| KNO<sub>3</sub> | 1% or 0.1M | 10 | 990 uM |

### Flow cytometry

*Physalis grisea* wild-type seeds were directly sown in soil in 96-cell plastic trays and grown to 4-week-old seedlings in the greenhouse. Two 1 cm disks from one leaf of wild type and 2 different regenerated plants were placed in separate Petri plates containing 0.5 mL of extraction buffer (CyStain UV Precise P, Partec, Münster, Germany) and chopped into small pieces using a razor blade. The suspension was filtered through a 50 μM filter. The suspension was analyzed using a flow cytometer (Becton Dickinson FACSAria II SORP Cell Sorter) and peak profile used to determine ploidy.

## Results and Discussion

### Ploidy determination of regenerated plants by pollen tube number

We explored the utility of germinating pollen and basing ploidy level determination on the number of pollen tubes [26]. This approach was attractive for our work compared to cytological methods, which can be time consuming and tedious when chromosomes are difficult to discern [1]. In addition, the pollen tube method had the potential to be higher throughput for assessment compared to other reported methods, would not require a high level of technical expertise, be less expensive, and did not require specialized equipment compared to other methods such as flow cytometry [1]. While the pollen germination method has advantages for distinguishing between diploid and polyploid status, flow cytometry, chromosome counts, or other methods would be needed to determine the exact ploidy.

Before assessing our experimental material for ploidy level, we tested the pollen germination method with leaf material from *P. grisea* wild-type and tissue culture regenerated plants by using flow cytometry confirmation (Fig. 1). For the 2 wild-type plants, the DNA content of each resulted in a peak representing nuclei at the G1 phase (channel 50) and another peak (PK2) of nuclei at G2 phase (channel 100) (Fig. 1A, B). These results indicated the lines were diploid. Germinated pollen from wild-type plants had only 1 tube, therefore, this confirmed 1 pollen tube indicated diploid. We then looked at the DNA content of 3 regenerated lines that were suspected of being polyploid based on phenotype. One plant was determined to be triploid based on flow cytometry results (Fig 1C) and the two remaining lines were determined to be tetraploid (Fig. 1E, D).

**Figure 1:**
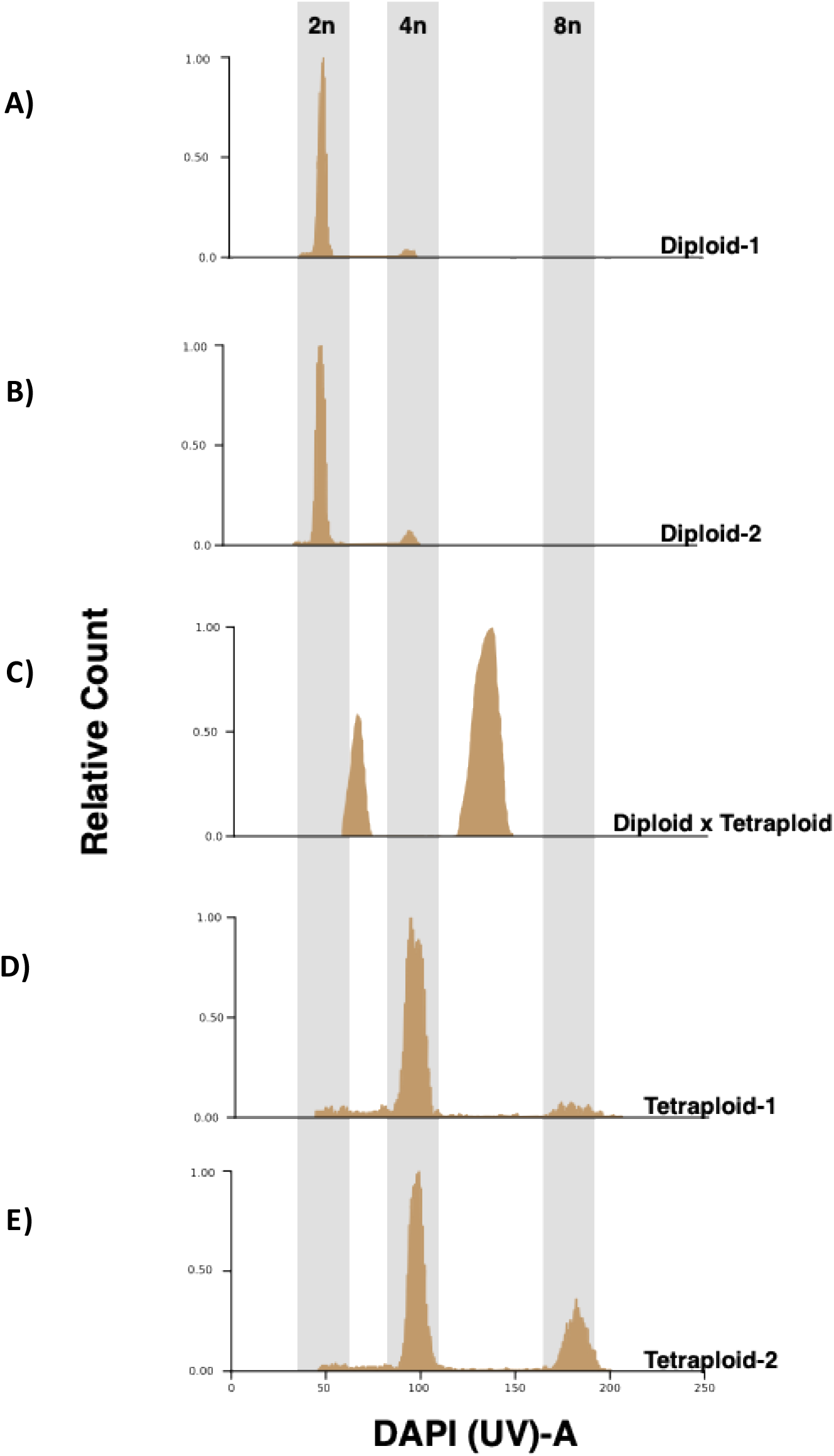
DNA ploidy analysis by flow cytometry of *Physalis grisea* wild-type and tissue culture regenerated plants. A, B: Histogram of DNA content of diploid wild-type plants showing a peak representing nuclei at G1 phase (channel 50) and another peak (PK2) of nuclei at G2 phase (channel 100); C: Histogram of a regenerated triploid plant with PK1 (channel 75) and PK2 (130) of nuclei at G1 and G2 phases; D, E: Histogram of a regenerated tetraploid plant with PK1 (channel 100) and PK2 (170) of nuclei at G1 and G2 phases.

For determining the ploidy status of regenerants, we selected 10 plants recovered from each medium treatment and explant origin except for *P. grisea* hypocotyl explants cultured on medium containing 0.25 and 0.5 mg/l zeatin where only 2 and 4 regenerants were recovered, respectively (Table 2). This lower number of regenerants is because we found the optimal concentrations for efficient *P. grisea* plant regeneration are 1 and 2 mg/l zeatin. Regenerated plants from all species were transferred to soil, grown to maturity in a greenhouse, and pollen was collected from flowers at the first sign of dehiscence. To obtain sufficient pollen for testing, the following number of flowers were collected for each species, 2 – 3 for *P. grisea*, 1 for *S. lycopersicum*, and 1 – 2 for *S. prinophyllum*. The incubation time in PGM depended on pollen tube development, therefore, tube number was counted over the course of 2 – 4 hrs. If 1 pollen tube was observed per pollen grain (Fig. 2), the origin plant was designated as diploid based on flow cytometry results with wild-type plants (Fig.1 A, B), whereas 2 tubes indicated a polyploid plant (Fig. 2). Considering all the zeatin concentrations, explant types, and species, 0 to 81% of the regenerated plants were found to be polypoid (Table 2).

**Table 2:** Percent of polyploid plants regenerated from seedling-derived explants cultured on plant regeneration medium containing various levels of zeatin.

| Species | Explant source | Zeatin concentration (mg/l) | Number of plants | Number of polyploid plants | % polyploid plants |
| --- | --- | --- | --- | --- | --- |
| *Physalis grisea* | Hypocotyl | 2 transfer to 1<sup>b</sup> | 10 | 8 | 80 |
|  |  | 1 | 10 | 8 | 80 |
|  |  | 0.5 | 4 | 3 | 75 |
|  |  | 0.25 | 2 | 2 | 100 |
|  | Cotyledon<sup>a</sup> | - | - | - | - |
| *Solanum lycopersicum* | Hypocotyl | 2 transfer to 1 | 10 | 8 | 80 |
|  |  | 1 | 10 | 3 | 30 |
|  |  | 0.5 | 10 | 4 | 40 |
|  |  | 0.25 | 10 | 2 | 20 |
|  | Cotyledon | 2 transfer to 1 | 10 | 1 | 10 |
|  |  | 1 | 10 | 0 | 0 |
|  |  | 0.5 | 10 | 0 | 0 |
|  |  | 0.25 | 10 | 2 | 20 |
| *Solanum prinophyllum* | Hypocotyl | 2 transfer to 1 | 10 | 2 | 20 |
|  |  | 1 | 10 | 3 | 30 |
|  |  | 0.5 | 10 | 5 | 50 |
|  |  | 0.25 | 10 | 4 | 40 |
|  | Cotyledon | 2 transfer to 1 | 10 | 4 | 40 |
|  |  | 1 | 10 | 3 | 30 |
|  |  | 0.5 | 10 | 0 | 0 |
|  |  | 0.25 | 10 | 1 | 10 |

<sup>a</sup>*Physalis grisea* cotyledon-derived explants do not regenerate plants [23].
<sup>b</sup>Transfer of explants from an initial plant regeneration medium containing 2 mg/l zeatin to 1 mg/l zeatin after 2 weeks according to previous reports [22, 23]. This is the standard method followed for plant regeneration from each species.

**Figure 2.**
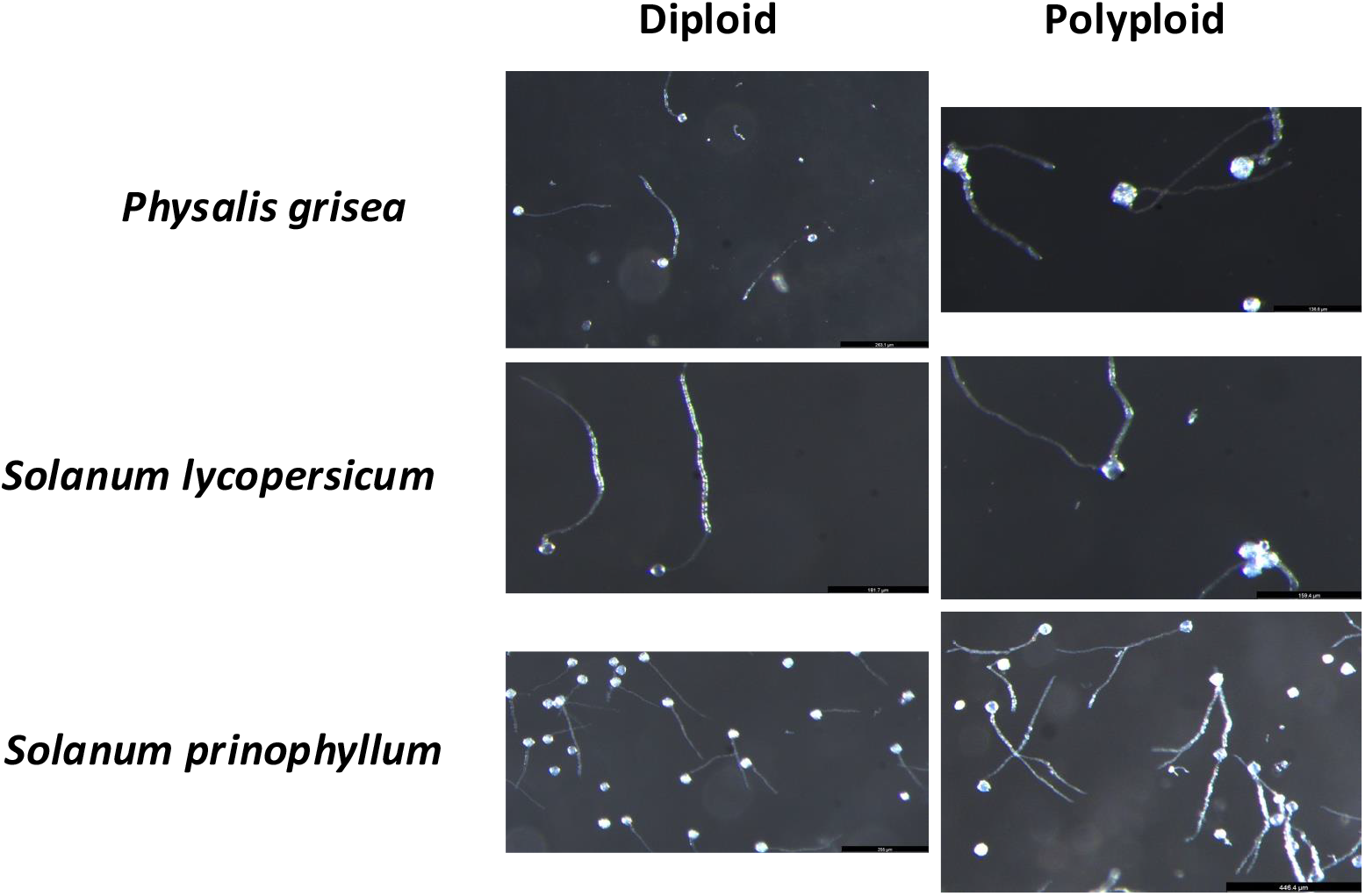
Determination of ploidy level based on the number of tubes on germinated pollen from plant regenerants. Germinated pollen with one tube indicates a plant is diploid, two tubes indicates it is polyploid.

### Effect of zeatin concentration on ploidy level

To investigate the effect of zeatin concentration on ploidy level of regenerated plants, we followed our standard method reported for *P. grisea* and *S. lycopersicum* where the initial plant regeneration medium contains 2 mg/l zeatin for 2 weeks of culture. However, for subsequent bi-weekly transfers, the medium contains only 1 mg/l zeatin [22, 23]. This step down from 2 to 1 mg/l was found to promote shoot elongation compared to constant culture on medium containing 2 mg/l zeatin. In this study, the same methods were applied to *S. prinophyllum* based on previous work with this species (unpublished results).

For *P. grisea*, there were no major differences observed in the percent of plants (75 – 100 %) regenerated on the different zeatin concentrations (Table 2). Although, considering that 0.25 and 0.5 mg/l are not optimum concentrations for *P. grisea* plant regeneration, only 2 and 4 plants, respectively, were recovered resulting in a small sample size.

The standard method for *S. lycopersicum* resulted in the largest number of polyploid plants at 80% from hypocotyls with a downward trend as the zeatin concentration decreased with only 20% plants recovered on medium containing 0.25 mg/l. However, for regenerants from cotyledons we did not observe a similar trend. The percent of polyploid plants was significantly lower at 10% on the standard method with no polyploid pants recovered on 0.5 and 1, but 20% on medium containing 0.25 mg/l. These results indicate that perhaps zeatin concentration has more influence on polyploidization in hypocotyls compared to cotyledons.

As for *S. prinophyllum*, we did not observe a trend of a higher incidence of polyploidization of plants regenerated as zeatin concentration increased. It was the opposite that as the concentration decreased the incidence increased. We hypothesize that perhaps a higher zeatin concentration suppresses endoreduplication in *S. prinophyllum* hypocotyl cells. However, like *S. lycopersicum*, there was a decreasing trend of polyploid regenerants recovered from cotyledon explants with 40% from the standard method and only 0 and 10% on 0.5 and 0.25 mg/l, respectively.

To our knowledge, there are no reports of the effects of zeatin-containing plant regeneration medium on the ploidy status of regenerated plants. Although, plant growth regulators, primarily auxins and cytokinins, have been reported to influence ploidy level of plants regenerated through effects on the mitotic cycle in the cells of explants [2]. For example, a plant growth regulator effect on the ploidy status of cells in *Asparagus officinalis* stem explants was shown to be affected by growth regulators in the medium with the auxin, 1-naphthaleneacetic acid, having the strongest effect [6].

### Effect of explant type on ploidy level

To determine if explant source contributed to the occurrence of polyploidization, we chose cotyledons and hypocotyls from in vitro-grown seedlings because they result in efficient plant regeneration from our species of interest [22, 23]. As indicated earlier, we found that *P. grisea* cotyledon explants did not regenerate shoots according to our standard protocol, therefore, only hypocotyls were cultured for this study [23]. The percent (81) of regenerated shoots found to be polyploid across all zeatin concentrations was significantly greater than those regenerated from hypocotyl explants of *S. lycopersicum* and *S. prinophyllum* at 43% and 35%, respectively (Table 3). In comparison with plants regenerated from cotyledon explants, 8% of S. *lycopersicum* and 20% of *S. prinophyllum* were polyploid. The lower percentages of polyploid regenerants compared to those regenerated from hypocotyl explants indicates that for future experiments the explant source of choice should be cotyledons.

**Table 3:** Collective number of polyploid plants regenerated from hypocotyl- and cotyledon-derived explants cultured on medium containing various zeatin concentrations.

| Species | Explant source | Number of plants | Number of polyploid plants | % polyploid plants |
| --- | --- | --- | --- | --- |
| *Physalis grisea* | Hypocotyl | 26 | 21 | 81 |
|  | Cotyledon<sup>a</sup> | - | - | - |
| *Solanum lycopersicum* | Hypocotyl | 40 | 17 | 43 |
|  | Cotyledon | 40 | 3 | 8 |
| *Solanum prinophyllum* | Hypocotyl | 40 | 14 | 35 |
|  | Cotyledon | 40 | 8 | 20 |

<sup>a</sup>*Physalis grisea* cotyledon-derived explants do not regenerate plants [23].

The effect of explant source on the polyploidization of regenerated plants has been previously reported for *S. lycopersicum* [6, 7, 10]. In the report by van den Bulk et al. (1990), they looked at ploidy variation in plants regenerated from leaf, cotyledon, and hypocotyl explants for *S. lycopersicum* cv Moneymaker [10]. The ploidy level observed was primarily tetraploid. For hypocotyl-derived explants, 50% of regenerants were polyploid, whereas only 1.5% and 12% from leaf and cotyledon explants, respectively, were polyploid. To determine the possible cause for these differences, they analyzed the ploidy status of the cells in the explants before culturing on plant regeneration medium. The results showed that hypocotyl-derived explants contained a mixture of diploid and polyploid cells, a condition referred to as polysomaty. In comparison, leaf- and cotyledon-derived explants contained either no or a lower population of polyploid cells.

A polysomaty condition in explants has also been reported for other species including *Solanum melongena* (eggplant) and *A*. officinalis [6, 27, 28]. For *S. melongena* hypocotyl, cotyledon, and leaf-derived explants, a similar trend was reported for the polysomaty condition as was for *S. lycopersicum* cv Moneymaker with leaf-derived explants having a lower polysomaty condition where no tetraploid cells were observed [10, 27]. However, leaf explants of *S. melongena* did not result in efficient plant regeneration as compared to cotyledon and hypocotyl explants. Between 25 to 50% of the plants regenerated from these explants were tetraploid. For *A. officinalis*, internodal stem explants were assessed for polysomaty and it was found that as the distance increased from the shoot apex, the percentage of polyploid cells also increased from predominantly 2C cells to 4C and 8C. The ploidy level of regenerated *A. officinalis* plants parallelled the polysomatic condition of the explant source [6].

Although the ploidy condition of explants from some species has been implicated, it is not always a predictor as was shown for plant regeneration from *Cannabis sativa* and *Humulus lupulus* (common hops) [28, 29]. *C. sativa* cotyledon, hypocotyl and leaf explants contained a mix of diploid and triploid cells; however, this did not strongly influence the ploidy status of regenerated plants where all regenerated plants were diploid. Although, a small percentage were found to be mixaploid meaning they contained diploid and tetraploid cells. For *H. lupulus*, all explant material (internodes, petioles, leaf discs) was diploid, but plants regenerated were tetraploid [11].

For some plant regeneration methods, there is an indirect production of plants through callus that develops from the explant. Callus is a rapidly dividing, undifferentiated mass of cells that when provided with exogenous growth regulators (phytohormones), namely cytokinins, for plant regeneration, the cells differentiate into plants. The callus phase can contribute to polyploid plants as was shown for *Dianthus acicularis* [12]. Polysomaty was not observed in the *D. acicularis* leaf explants, but was detected in the calli that developed [12]. Considering that *P. grisea* hypocotyl explants produce more callus prior to the initiation of plant regeneration compared to *S. lycopersicum* and *S. prinophyllum*, this points to a potential cause of the high percentage of polyploid regenerants.

While we did not assess the polysomatic condition of hypocotyls and cotyledons from our species of interest, based on the reports for *S. lycopersicum* and eggplant, we can predict from our results that hypocotyls possibly have a higher percentage of polyploid cells compared to cotyledons [10, 27]. Although, it is possible that if there was a mixture of diploid and polyploid cells, perhaps the polyploid cells had a fitness advantage over diploid cells for plant regeneration. It is possible that since we selected the first 10, fully developed plants that regenerated, this could have skewed plant recovery in favor of polyploid regenerants.

### Effect of plant species

For plants regenerated from hypocotyls from our 3 species of interest across all zeatin concentrations, we found that 81% of *P. grisea* plants were polyploid compared to 43% and 35% for *S. lycopersicum* and *S. prinophyllum*, respectively (Table 3). As for ploidy level of plants regenerated from cotyledons, 8% of *S. lycopersicon* and 20% of *S. prinophyllum* plants were polyploid. Based on these results, it appears that in addition to explant type, there might also be a species effect on the recovery of polyploid regenerants.

In a report by Ellul et al. (2003) on *Agrobacterium tumefaciens*-mediated transformation of *S. lycopersicum*, they observed a difference in ploidy level of regenerants using the same method across 5 different genotypes. The percent of tetraploid plants recovered ranged from 20 to 57%. They indicated that the higher ploidy levels were not correlated to the proportion of 4C and 8C nuclei in the cotyledon explants leaving the possibility of an intrinsic interaction of the genotype with a factor in either the plant regeneration or transformation process. A genotype effect on ploidy was also observed for *H. lupulus* plant regenerants where one genotype had a high frequency of tetraploid plants in comparison to 2 other genotypes [29].

### Effect of ploidy level on phenotypic variation

Plant regenerants determined to be polyploid based on the pollen germination test were transferred to the greenhouse and grown to maturity for phenotypic assessment compared to their wild-type counterparts. For *P. grisea* and *S. prinophyllum*, 2 diploid and 2 polyploid plants per species were transferred to the greenhouse. However, for *S*. lycopersicum we transferred 10 diploid and 10 polyploid plants to the greenhouse because we found that the polyploid plants took longer to flower. All plants were evaluated for differences in growth habit, flowers, fruit, and seeds compared to their wild-type, diploid counterparts (Figs. 3 – 5). In general, there were noticeable increases in flower and seed size in polyploid regenerants across all 3 species. Whereas for fruit, there was only a slight increase in size for *S. lycopersicum* (Fig. 4) and no noticeable significant differences for either *P. grisea* or *S. prinophyllum*. There were also no significant differences in plant size for any of the species.

**Figure 3.**
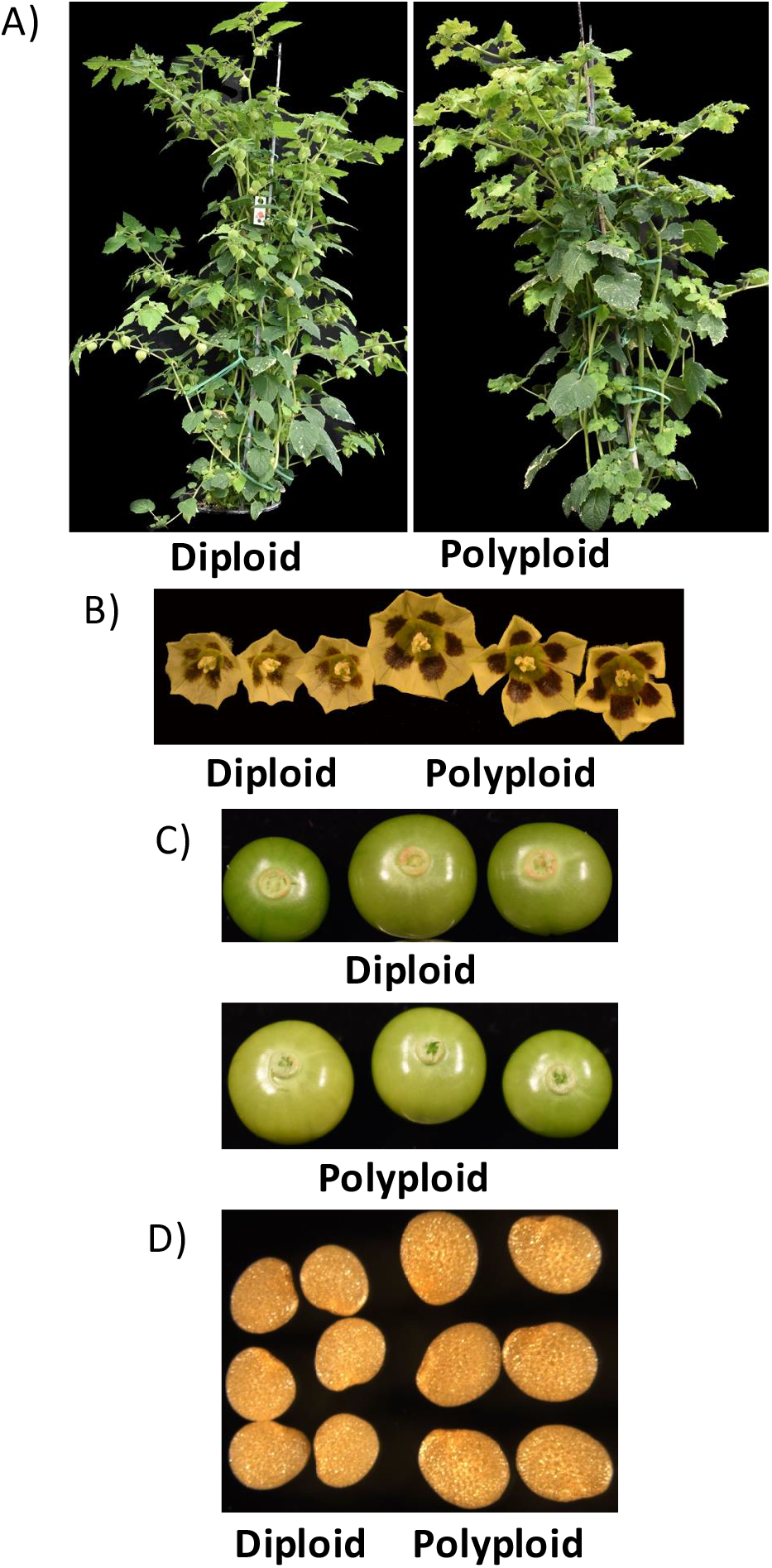
Phenotypes of diploid and polyploid *Physalis grisea* plant regenerants. A: Mature, greenhouse-grown plants; B: Flowers; C: Fruit; D: Seeds

**Figure 4.**
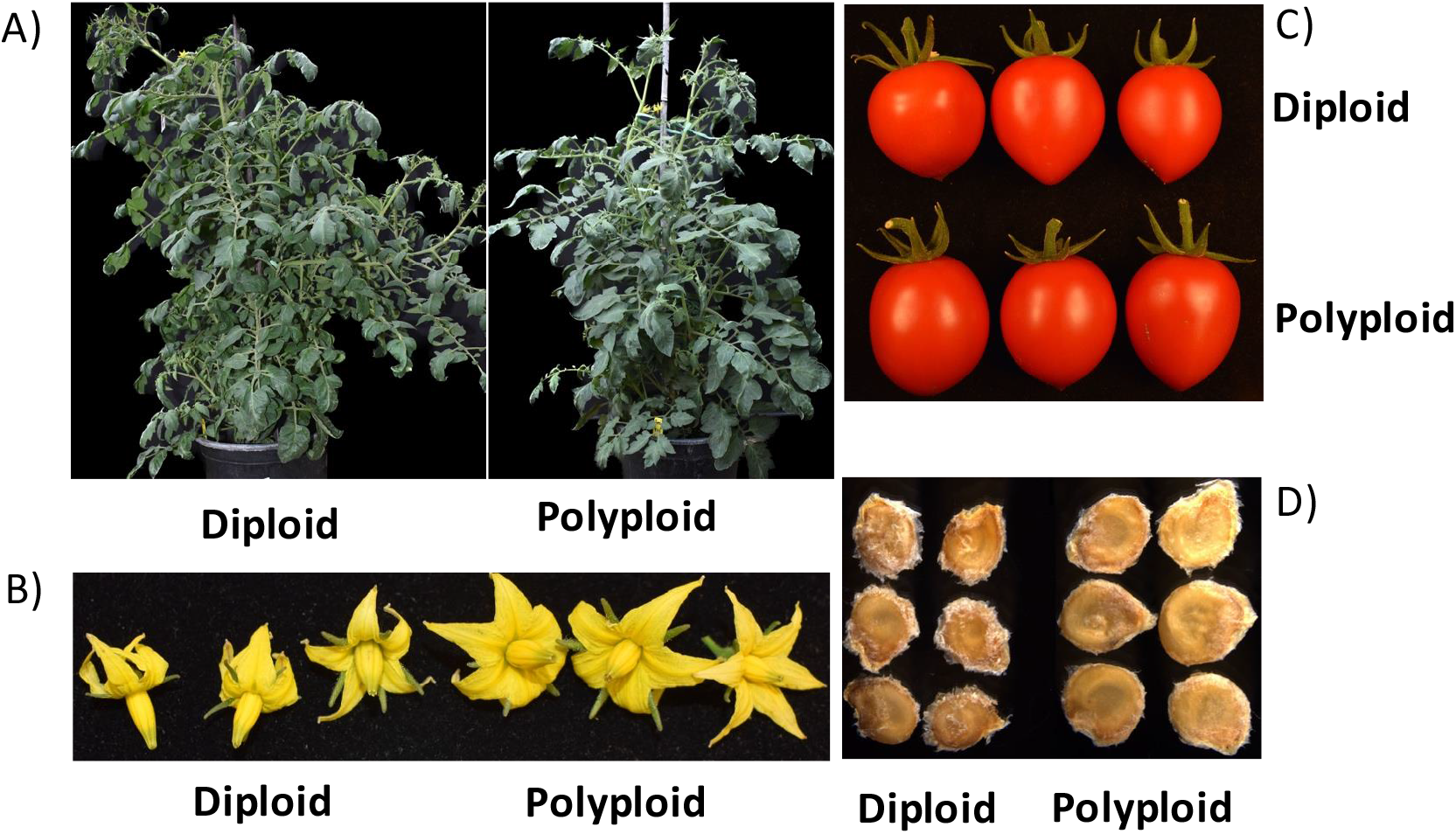
Phenotypes of diploid and polyploid *Solanum lycopersicum* plant regenerants. A: Mature, greenhouse-grown plants; B: Flowers; C: Fruit; D: Seeds

**Figure 5.**
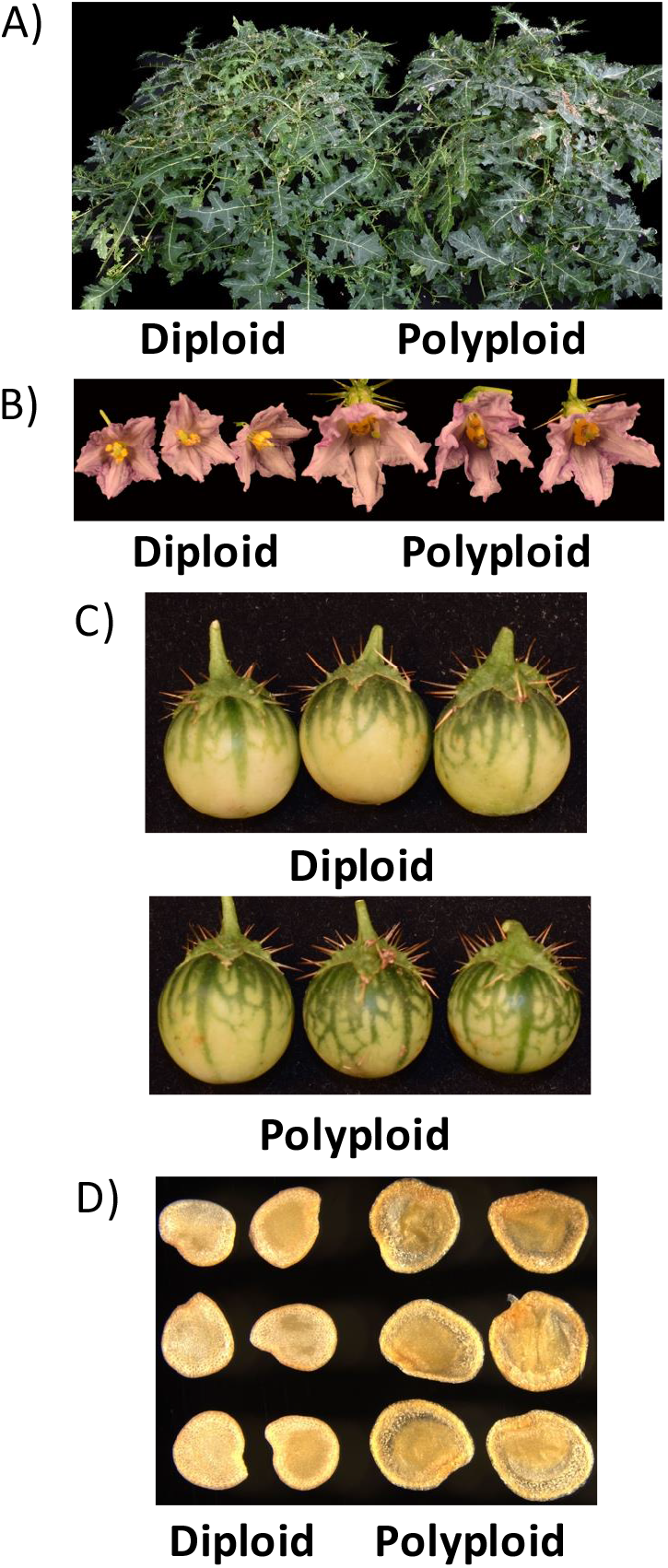
Phenotypes of diploid and polyploid *Solanum prinophyllum* plant regenerants. A: Mature, greenhouse-grown plants; B: Flowers; C: Fruit; D: Seeds

While a deviation from a normal chromosome number has been shown to impact plant morphology, it is at times not a direct correlation for all species or for overall development [3]. In *Cucumis melo* (honeydew melon), it was shown that polyploid regenerants did not exhibit any differences in morphology compared to diploid [8]. And as was shown in tamarillo regenerants, chromosome number is not always a predictor of phenotype because even diploid plants showed abnormalities, which points to other possible sources of variation, such as epigenetic changes that can arise from plant tissue culture conditions [30]. In some cases, an increase from diploid to tetraploid in regenerants is positively viewed when the change in ploidy results in for example, an increase in fruit size making the material useful for plant breeding programs [10]. Therefore, while morphological changes are not desirable in some applications, there is value in the crop improvement of different species especially ornamentals where chromosome doubling is used in breeding programs [31]. For breeding of triploid *H. lupulus*, diploid lines need to be brought to the tetraploid level. There was interest in exploring plant tissue culture to generate tetraploid lines instead of applying antimitotic compounds for chromosome doubling, which increases the frequency of chimeric plants where the ploidy level is unstable when vegetative micropropagation is used to increase the number of plants [29].

## Conclusion

The occurrence of ploidy changes in plants derived from in vitro culture has both positive and negative implications. On the positive side, polyploid plants often exhibit desirable traits such as increased size, enhanced stress resistance, and improved yield, making them valuable for agriculture [32]. Conversely, unexpected ploidy changes can lead to undesirable characteristics such as reduced fertility, abnormal growth, and genetic instability. Therefore, identifying the source and managing ploidy status during tissue culture is essential for ensuring the production of high-quality plants. As outlined in this report, careful consideration of various tissue culture parameters, including explant type and plant growth regulators, in parallel with regular monitoring for ploidy variation are critical for obtaining desirable outcomes.

## Acknowledgements

This study was supported by funding awarded to J.V.E. and Z.B.L. from the National Science Foundation Plant Genome Research Program grant IOS-2216612. K.S and Y.G would like to thank Elise Tomaszewski for her consultation on the pollen germination protocol.

## Author contribution statement

K.S., Y.M., and J.V.E. designed experiments; K.S. and Y.M. performed experiments. I.G. performed the flow cytometry and provided the pollen germination protocol. J.V.E wrote the manuscript.

